# PRIorI: a graph-based mining of structural arrangements in protein-protein interfaces

**DOI:** 10.64898/2025.12.03.692053

**Authors:** Vagner Soares Ribeiro, Vinicius de Almeida Paiva, Isabela de Souza Gomes, Lizandra Morgado Santos, Hélio Henrique Medeiros Silva, Vinicius Vitor dos Santos Dias, Sabrina de Azevedo Silveira

## Abstract

**Summary:** Protein-protein interactions (PPIs) are fundamental to biological processes and central to understanding disease mechanisms, making them essential for drug discovery, peptide-based therapeutics, and vaccine development. Identifying conserved structural arrangements within PPI interfaces can provide valuable insights into molecular recognition and interaction mechanisms. Here, we introduce PRIorI (PRotein-PRotein InteractiOn gRaph Isomorphism), a web-based platform to explore precomputed protein-protein interaction networks and identify conserved atomic-level interaction motifs within structural complexes. Unlike traditional methods, PRIorI models PPI interfaces as bipartite graphs, applying graph isomorphism techniques to efficiently retrieve interaction patterns independent of sequence alignment or structural superimposition. Users can query precomputed PDB interaction graphs, upload custom protein structures, or design structural motifs for targeted searches. To illustrate its applicability, we used PRIorI to analyze the SARS-CoV-2 Spike-ACE2 interface, identifying key interaction motifs previously reported in the literature. The platform successfully identified salt bridges, hydrogen bonds, and hydrophobic interactions that stabilize the viral-host complex, demonstrating PRIorI’s utility in protein interaction analysis.

**Availability and implementation:** Webserver implemented using typescript, python, Next.js and Express.js. Uses MongoDB for storage, Redis for cache and is hosted by Apache on an Ubuntu Server. PRIorI web platform is freely available at https://priori.ufv.br/.

**Supplementary information:** Supplementary data are available at *Bioinformatics* online.

## 1 Introduction

Proteins are fundamental macromolecules responsible for a wide range of biological processes, including structural support, enzymatic activity, and cellular regulation (Kessel and Ben-Tal, 2018; Nelson and Cox, 2017). Their interactions with other biomolecules are essential for maintaining homeostasis and orchestrating complex cellular functions (Zhong *et al*., 2019). Among these, protein-protein interactions (PPIs) are crucial in cell signaling, immune responses, and metabolic pathways (Jones and Thornton, 1996; Rao *et al*., 2014). Dysregulation of PPIs is associated with various diseases, such as cancer, neurodegenerative disorders, and infectious diseases, making them critical targets for therapeutic interventions (Richards *et al*., 2021; Greenblatt *et al*., 2024; Safari-Alighiarloo *et al*., 2014).

With the continuous expansion of protein structure databases, resources such as the Protein Data Bank (PDB) (Berman *et al*., 2000) have become indispensable for characterizing protein domains and interaction interfaces. The PDB has grown significantly and currently contains over 230,000 experimentally determined structures, providing a rich dataset for structural biology research. However, despite this extensive collection, many proteins remain functionally uncharacterised. While the 2021 Pfam release reported 4,244 families classified as domains of unknown function (DUFs), corresponding to approximately 23% of all families, more recent data indicate that Pfam contains 4,795 DUF-designated families, and that only 7,769 of the 19,632 families belong to a clan (Ponamareva *et al*., 2024). Additionally, recent advances in AlphaFold (Jumper *et al*., 2021; Varadi *et al*., 2024) have led to the prediction of complete proteomes with near-experimental accuracy, further expanding the available structural data and creating new opportunities for functional annotation, drug discovery, and the development of targeted therapies.

Several computational approaches have been developed to analyze protein interactions and structural motifs. Proteins Plus is a web-based platform that provides a growing collection of molecular modeling tools, enabling users to perform structural analyses, molecular docking, and search for structural variants in protein and nucleic acid models (Schöning-Stierand *et al*., 2022). SiteMine enables large-scale binding site similarity searches across protein structure databases, focusing primarily on ligand interactions (Reim *et al*., 2024). GeoMine is a search engine for ligand-bound and predicted empty binding sites in the PDB, offering textual, numerical, and geometric search functionalities for user-defined interaction patterns (Diedrich *et al*., 2021, 2024). GSP4PDB is a web-based system for searching structural patterns in proteins, optimized through denormalization and indexing to improve query performance for graph-based structural motifs (Angles *et al*., 2024). Arpeggio calculates atomic-level non-covalent interactions within protein structures (Jubb *et al*., 2017), while HyPPI (Schöning-Stierand *et al*., 2022) aids in predicting functional PPI hotspots. CrossMiner enables pharmacophore-based searches, but its flexibility is constrained by predefined descriptors (Korb *et al*., 2016). However, few tools support user-defined searches for conserved structural arrangements in protein-protein interfaces, limiting researchers’ ability to explore interaction patterns intuitively and in a customizable manner.

To address this gap, we introduce PRIorI (PRotein-PRotein InteractiOn gRaph Isomorphism), a web-based platform designed to analyze and explore precomputed protein-protein interaction interfaces at the atomic level. PRIorI models PPI interfaces as bipartite graphs and applies graph isomorphism techniques to efficiently retrieve interaction networks within structural complexes. Unlike previous approaches, it allows users to investigate conserved interaction motifs directly within known protein complexes, without relying on sequence alignment or structural superimposition. The methodology behind PRIorI is based on ppiGReMLIN (Queiroz *et al*., 2020), which applies graph-based mining techniques to detect conserved structural arrangements in protein-protein interfaces, identifying recurring interaction patterns at the atomic level. PRIorI provides a valuable resource for characterizing PPI interfaces, generating structural hypotheses, and supporting research in structural biology, drug discovery, and protein engineering by enabling the exploration of potential ligand interactions and the identification of novel therapeutic targets through an intuitive interface and an efficient graph-based approach.

## 2 Platform description

PRIorI is a web-based tool that analyzes and explores precomputed interaction networks within protein-protein complexes. It enables users to examine atomic-level interaction patterns in structural complexes stored in the Protein Data Bank (PDB). The system provides three modes for interaction exploration: database search by clicking on *Explore Database*, custom PDB structure submission by clicking on *Submit PDB*, and manual motif design by clicking on *Graph Input*.

In *Explore Database*, users can input a PDB ID and specify interacting chains to retrieve precomputed interaction graphs representing the atomic contacts between chains. These graphs are dynamically retrieved from a MongoDB database using PyMongo queries, ensuring efficient access to stored interaction networks without requiring persistent user-specific storage.

Alternatively, PRIorI allows users to upload a custom PDB file in *Submit PDB*, enabling the system to compute the interface graphs for a user-provided protein complex. Once processed, the results are stored, and the user is notified via email when they are ready for visualization. This functionality (*custom PDB structure submission*) enables researchers to analyze interaction networks within newly modeled or experimentally solved protein complexes.

The third mode to explore PPI interactions with PRIorI allows users to manually define a structural query by drawing an interaction motif. Users specify atoms, their physicochemical properties, and interactions in this mode to build a customized interface graph. Once submitted in *Graph Input*, PRIorI compares the designed motif to precomputed PPI interaction graphs for the whole PDB, helping users explore whether a specific structural arrangement appears in known complexes. This approach is particularly useful for studying conserved molecular recognition patterns and assessing the structural feasibility of engineered interactions.

PRIorI models protein-protein interfaces as bipartite graphs, where nodes represent atoms and edges represent interactions between atoms from different chains. The dataset used for graph generation was obtained from the PDB on May 14, 2025, and includes only entries containing protein chains, totaling 90,184 structures. Interaction graphs were computed using an adapted version of the ppiGReMLIN algorithm, which applies a cutoff-based strategy to define atomic interactions. The system generated 4,354,271 bipartite graphs in JSON format within a MongoDB 4.4 database. Using a non-relational database ensures efficient storage and rapid querying, handling variability in graph size while allowing for asynchronous query execution. Users receive a notification link once their analysis is complete. PRIorI is freely accessible and does not require user registration or login.

Upon query completion, PRIorI presents results in a structured format, listing interacting atoms, their respective chains, and direct links for visualization. The three-dimensional molecular viewer, powered by NGL Viewer, provides an interactive interface representation, allowing users to rotate, zoom, and modify visual settings for detailed structural analysis. Additionally, PRIorI features a graph-based visualization using Python’s Bokeh library, offering an alternative exploration of interface graphs. The graph visualization is color-matched to the molecular viewer, and users can zoom, pan, and hover over nodes and edges to inspect atomic-level details, facilitating deeper structural interpretation.

PRIorI introduces a powerful capability by allowing users to analyze interaction patterns within a protein complex and compare them with precomputed interface interaction data. This functionality supports the identification of conserved interaction motifs within PPIs, enabling detailed structural exploration, refinement of molecular hypotheses, and investigation of interaction modifications. These insights have broad applications in structural biology, protein engineering, and drug discovery, where understanding recurring interface interaction networks is crucial for designing inhibitors, optimizing protein interfaces, and elucidating protein function. Figure 1 presents a workflow summarizing PRIorI’s functionalities, demonstrating its application in structural motif analysis.

**Fig. 1.**
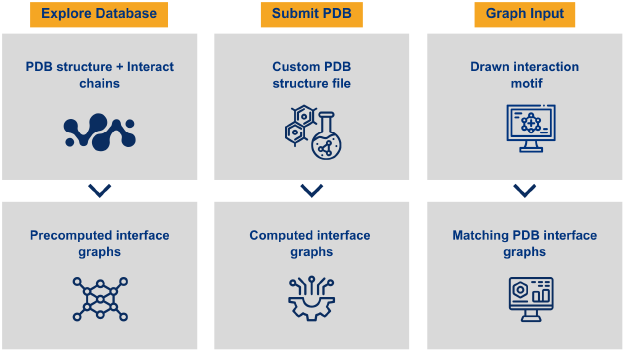
PRIorI workflow and functionalities. PRIorI provides three distinct modes to explore protein-protein interaction interfaces: (i) Explore Database, where users input a PDB ID and chains to access precomputed interface interaction graphs; (ii) Submit PDB, allowing users to upload their own protein structures for on-demand calculation of interface graphs; and (iii) Graph Input, where users manually define atomic-level interaction motifs to search for matching interface graphs in the PDB database.

## 3 Applications

The interaction between the SARS-CoV-2 Spike glycoprotein and the human ACE2 receptor is essential for viral entry into host cells. The Spike protein’s receptor-binding domain (RBD) establishes a high-affinity interface with ACE2 through a network of hydrogen bonds, salt bridges, and hydrophobic interactions, stabilizing the viral attachment process. Several structural studies (Shang *et al*., 2020; Yan *et al*., 2020; Lan *et al*., 2020) have identified key residues involved in this interaction, helping to elucidate the molecular mechanisms underlying SARS-CoV-2 inffectivity.

To evaluate how these interactions are represented in PRIorI, we queried the platform using the PDB structure 6M0J, which captures the high-resolution conformation of the Spike-ACE2 complex. PRIorI extracted and listed the atomic interactions at the interface, allowing us to compare them with those previously described in the literature. The results confirmed the presence of multiple interactions highlighted in previous structural studies. Among them, the Lys417 (Spike) – Asp30 (ACE2) salt bridge was detected, supporting its role in the electrostatic stabilization of the complex. Additionally, the hydrophobic cluster formed by Phe486 (Spike) interacting with Leu79, Met82, and Tyr83 (ACE2) was identified, reinforcing its importance in binding affinity. PRIorI also detected the Thr500 (Spike) – Tyr41 (ACE2) hydrogen bond, as well as a complex interaction network involving Gln449 (Spike) with Asp38 and Gln42 (ACE2). Furthermore, the results revealed additional stabilizing contacts, including Gly487 (Spike) engaging Gln24 and Tyr83 (ACE2), along with an interaction between Asn353 (ACE2) and Gly502 (Spike).

To further explore PRIorI’s capabilities, we used *Graph Input* to search for interaction patterns similar to the hydrophobic cluster formed by Phe486 (Spike) interacting with Leu79, Met82, and Tyr83 (ACE2) in structure 6M0J. This search identified several complexes with analogous interaction patterns, notably the structure 6M17. Within 6M17, significant interactions were observed involving residues Lys31 (ACE2) and Tyr489 (Spike) (Hatmal *et al*., 2020). Lys31 on ACE2 is frequently involved in stabilizing interactions, significantly contributing to the stability and formation of the complex by interacting closely with Spike protein residues (Pirolli *et al*., 2021). Tyr489 on the Spike protein is also critically important; its backbone is stabilized by hydrogen bonding, and its side chain engages in pi-pi aromatic interactions. This structural configuration enables Tyr489 to establish strong electrostatic interactions (Chowdhury *et al*., 2020). Identifying these residues further emphasizes their functional importance in the viral attachment process and highlights how PRIorI can effectively uncover conserved interaction motifs crucial for molecular recognition.

Additionally, to explore the potential side effects of therapeutic agents using PRIorI, we initially used the *database search mode*. We identified the residue Tyr505 (Spike) and its associated interactions, which are critical for binding efficacy. Tyr505 is essential in maintaining the structural integrity of the Spike-ACE2 complex, and its disruption can severely impair viral entry, making it a key target in drug discovery efforts (Lin *et al*., 2021). Leveraging this information, the *manual motif design mode* can be subsequently employed to explore analogous interaction patterns involving Tyr505 across other protein complexes within the PDB. Such an analysis enables the identification of off-target interactions that might explain clinical side effects observed in therapeutics targeting similar structural motifs.

Overall, PRIorI successfully identifies key interaction motifs in protein-protein interfaces, demonstrating its ability to characterize and compare complex interaction networks systematically. This makes PRIorI a valuable tool for researchers investigating molecular recognition mechanisms, drug design, and potential therapeutic targets. Two case studies are presented in the Supplementary Material to illustrate the potential of PRIorI in the study and characterisation of protein–protein interfaces, as well as in the detection of patterns within these interfaces and in the search for such patterns across the PDB.

## 4 Conclusion

PRIorI provides an efficient and flexible strategy for analyzing interface interaction networks within protein-protein complexes, enabling researchers to explore conserved atomic-level structural motifs in precomputed PPI interfaces. PRIorI offers a precise and adaptable framework for characterizing protein interactions by modeling PPIs as bipartite graphs and applying graph isomorphism techniques. Unlike residue-based approaches, PRIorI allows searches at the atomic level, capturing conserved interaction patterns with greater specificity without relying on sequence alignment or structural superimposition.

To illustrate its applicability, we used PRIorI to analyze the SARS-CoV-2 Spike-ACE2 complex (PDB: 6M0J). The tool successfully identified key hydrogen bonds, salt bridges, and hydrophobic interactions that stabilize the viral-host interface, aligning with structural data previously reported in the literature. By identifying these interactions, PRIorI demonstrated its potential for systematic analysis of protein interfaces, highlighting how it can be used to explore and interpret interface interactions and support structural hypothesis testing.

With an intuitive interface and interactive visualization capabilities, PRIorI facilitates large-scale structural motif analysis, making it a valuable resource for structural biology, drug discovery, and protein engineering. The platform is publicly available, efficiently processes queries, and returns results within minutes, allowing researchers to analyze and interpret molecular interactions rapidly. By enabling a user-driven approach to PPI exploration, PRIorI expands the possibilities for understanding molecular recognition, identifying novel therapeutic targets, and designing biomolecular interventions.

## Acknowledgements

Not applicable

