## Supplementary Material for "PRIorI: a graph-based mining of structural arrangements in protein-protein interfaces"

### Complementary Material of Study 1: Analysis of the 1HVR Complex in PRIorI

**Figure 1 - Structural representation of the complex between the RBD of the SARS-CoV-2 Spike protein and the human ACE2, analyzed using the PRIORI platform**

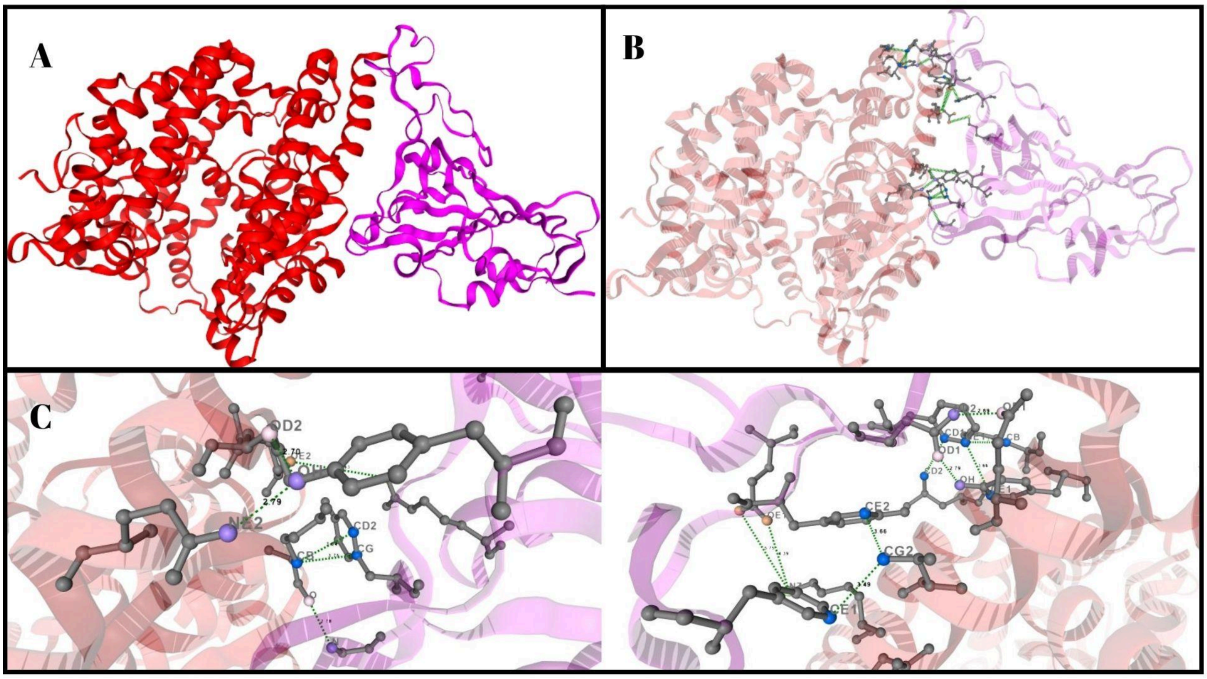


A) Overall three-dimensional structure of the complex obtained from the crystallographic model deposited in the Protein Data Bank. B) Three-dimensional model processed by PRIORI, highlighting the molecular contact regions. C) Detailed view of specific interactions at the Spike–ACE2 interface, showing participating residues and interatomic distances in ångströms (Å).

**Figure 2 - Visualization of the molecular interactions between the SARS-CoV-2 Spike RBD and human ACE2 generated by PRIORI. Colors distinguish different chemical natures.**

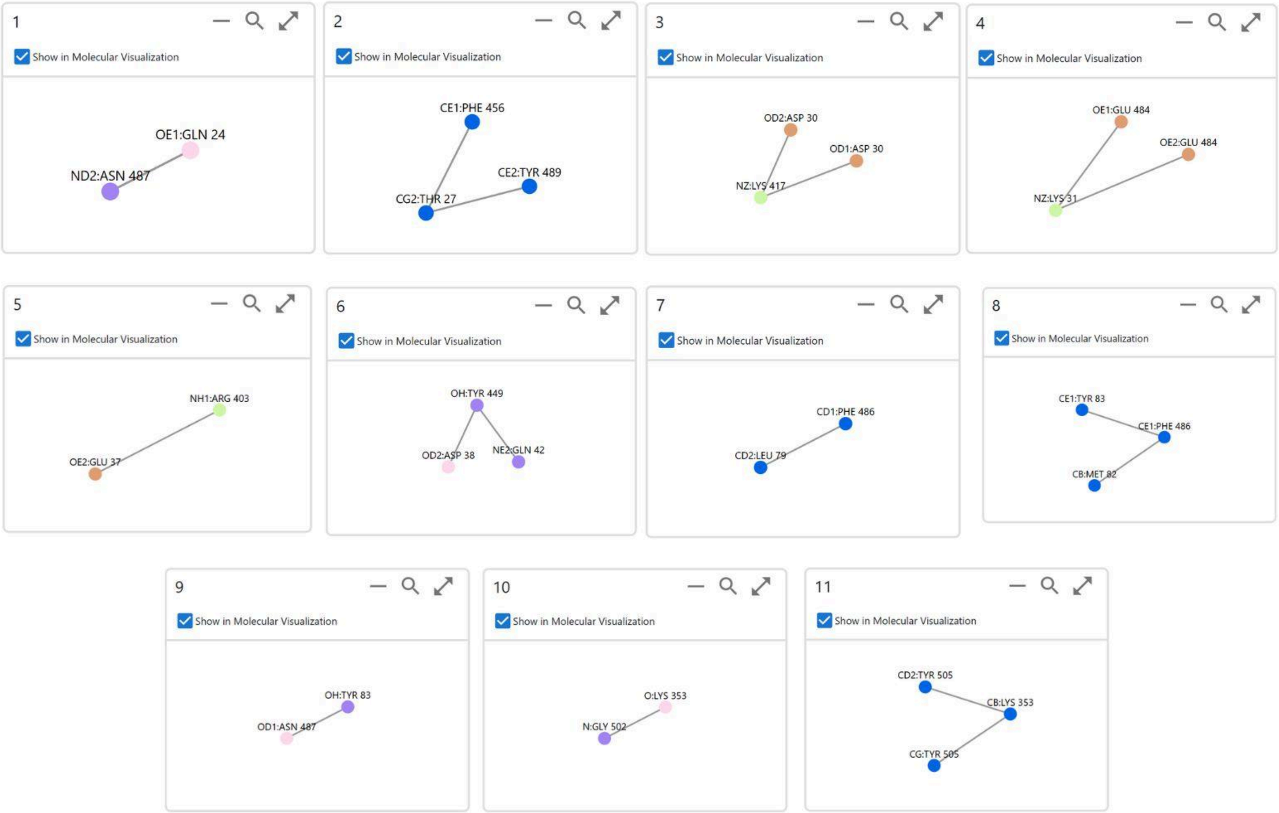

**Table 1 - Interaction profile between residues of the SARS-CoV-2 Spike RBD and human ACE2.**

| Index | Graph Number | Interaction Type | Atom 1 | Atom 2 | Distance |
|---|---|---|---|---|---|
| 1 | 1 | Hydrogen Bond | GLN-OE1-24-A (Acceptor) | ASN-ND2-487-E (Donor) | 2.69 Å |
| 2 | 2 | Hydrophobic | TYR-CE2-489-E (Hydrophobic) | THR-CG2-27-A (Hydrophobic) | 3.66 Å |
| 3 | 2 | Hydrophobic | THR-CG2-27-A (Hydrophobic) | PHE-CE1-456-E (Hydrophobic) | 3.49 Å |
| 4 | 3 | Salt Bridge | ASP-OD1-30-A (Negative) | LYS-NZ-417-E (Positive / Donor) | 4.03 Å |
| 6 | 4 | Salt Bridge | GLU-OE2-484-E (Negative) | LYS-NZ-31-A (Positive) | 5.75 Å |
| 7 | 4 | Salt Bridge | LYS-NZ-31-A (Positive) | GLU-OE1-484-E (Negative) | 4.39 Å |
| 8 | 5 | Salt Bridge | ARG-NH1-403-E (Positive) | GLU-OE2-37-A (Negative) | 5.63 Å |
| 9 | 6 | Hydrogen | GLN-NE2-42 | TYR-OH-449 | 2.79 Å |

| | | | | | |
|---|---|---|---|---|---|
| | | Bond | -A (Donor) | -E (Donor / Acceptor) | |
| 10 | 6 | Hydrogen Bond | ASP-OD2-38-A (Acceptor) | TYR-OH-449-E (Donor / Acceptor) | 2.70 Å |
| 11 | 7 | Hydrogen Bond | THR-OG1-500-E (Donor / Acceptor) | TYR-OH-41-A (Donor / Acceptor) | 2.71 Å |
| 12 | 8 | Hydrophobic | PHE-CD1-486-E (Hydrophobic) | LEU-CD2-79-A (Hydrophobic) | 3.77 Å |
| 13 | 9 | Hydrophobic | PHE-CE1-486-E (Hydrophobic) | MET-CB-82-A (Hydrophobic) | 3.58 Å |
| 14 | 9 | Hydrophobic | PHE-CE1-486-E (Hydrophobic) | TYR-CE1-83-A (Hydrophobic) | 3.66 Å |
| 15 | 10 | Hydrogen Bond | TYR-OH-83-A (Donor) | ASN-OD1-487-E (Acceptor) | 2.79 Å |
| 16 | 11 | Hydrogen Bond | LYS-O-353-A (Acceptor) | GLY-N-502-E (Donor) | 2.78 Å |
| 17 | 12 | Hydrophobic | LYS-CB-353-A (Hydrophobic) | TYR-CG-505-E (Hydrophobic) | 3.75 Å |
| 18 | 12 | Hydrophobic | LYS-CB-353-A (Hydrophobic) | TYR-CD2-505-E (Hydrophobic) | 3.64 Å |

**Complementary Case Study 2: Analysis of the 1HVR Complex in PRIorI**

The HIV-1 protease is a homodimeric aspartyl protease responsible for cleaving viral polyproteins, a process essential for viral maturation and infectivity. Inhibition of this enzyme prevents the formation of mature viral particles, making it one of the main therapeutic targets in HIV treatment. The crystallographic structure 1HVR, available in the Protein Data Bank, represents the HIV-1 protease in complex with an inhibitor, allowing direct observation of the molecular interactions that stabilize this complex.

**Figure 1 - Structural characterization of the HIV-1 protease–inhibitor complex (PDB: 1HVR).**

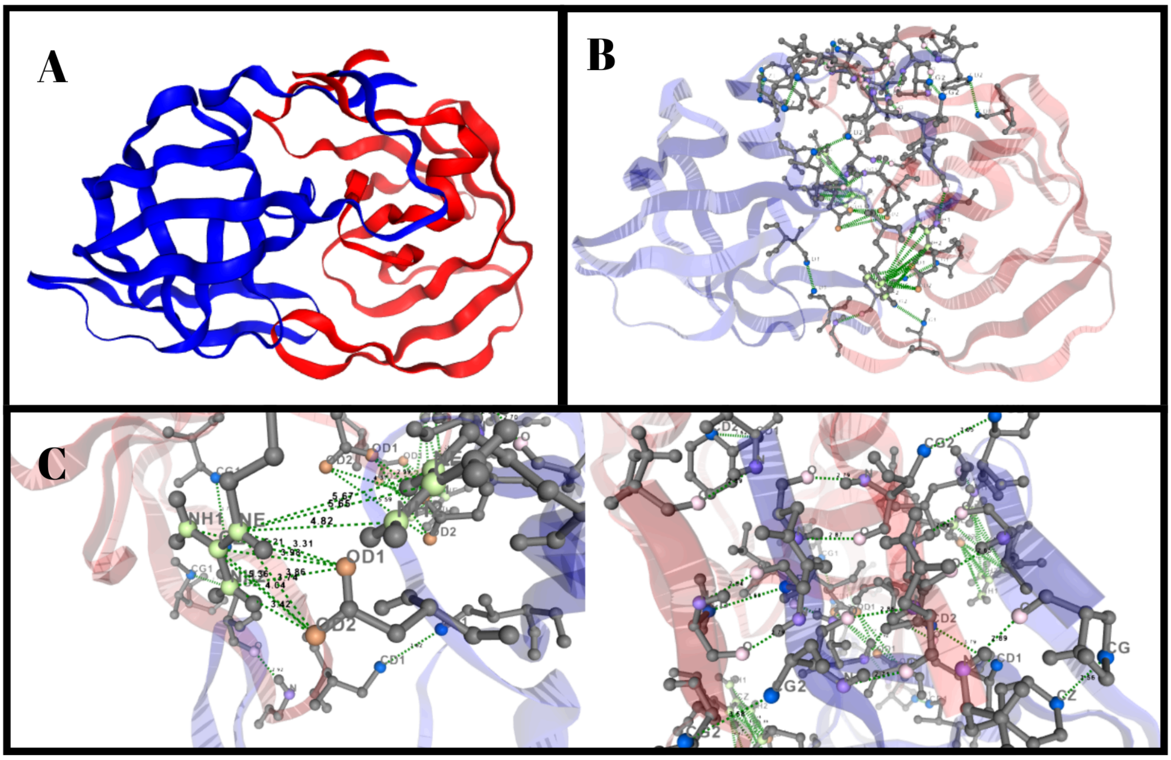


A) Overall structure of the HIV-1 protease in its homodimeric form, with the two monomers shown in blue and red. B) View of the protease–inhibitor complex, highlighting the position of the ligand within the catalytic cleft and its fit between the two monomers. C) Close-up of the non-covalent interactions identified using PRIorI, including hydrogen bonds and hydrophobic contacts between the inhibitor and key functional residues of the enzyme, with emphasis on the catalytic dyad and the flap regions.

Analysis of the structure using PRIorI identified the main non-covalent interactions between the catalytic and structural residues of the protease and the ligand. The results generated by the PRIorI highlighted hydrogen bonds and hydrophobic contacts involving residues Asp25 and Asp25′, which compose the enzyme's characteristic catalytic dyad. In addition, the system detected additional contacts with residues Gly27, Asp29, and Ile50 from both chains, forming a stabilizing network around the inhibitor that keeps it firmly positioned within the catalytic cleft.

The analysis also revealed significant hydrophobic interactions involving residues Pro81, Val82, and Ile84, which contribute to the stability and precise fit of the ligand in the active site. These contacts create an apolar environment complementary to the corresponding regions of the inhibitor, strengthening the affinity and specificity of binding.

**Figure 2 - Individual interaction graphs generated for the 1HVR complex.**

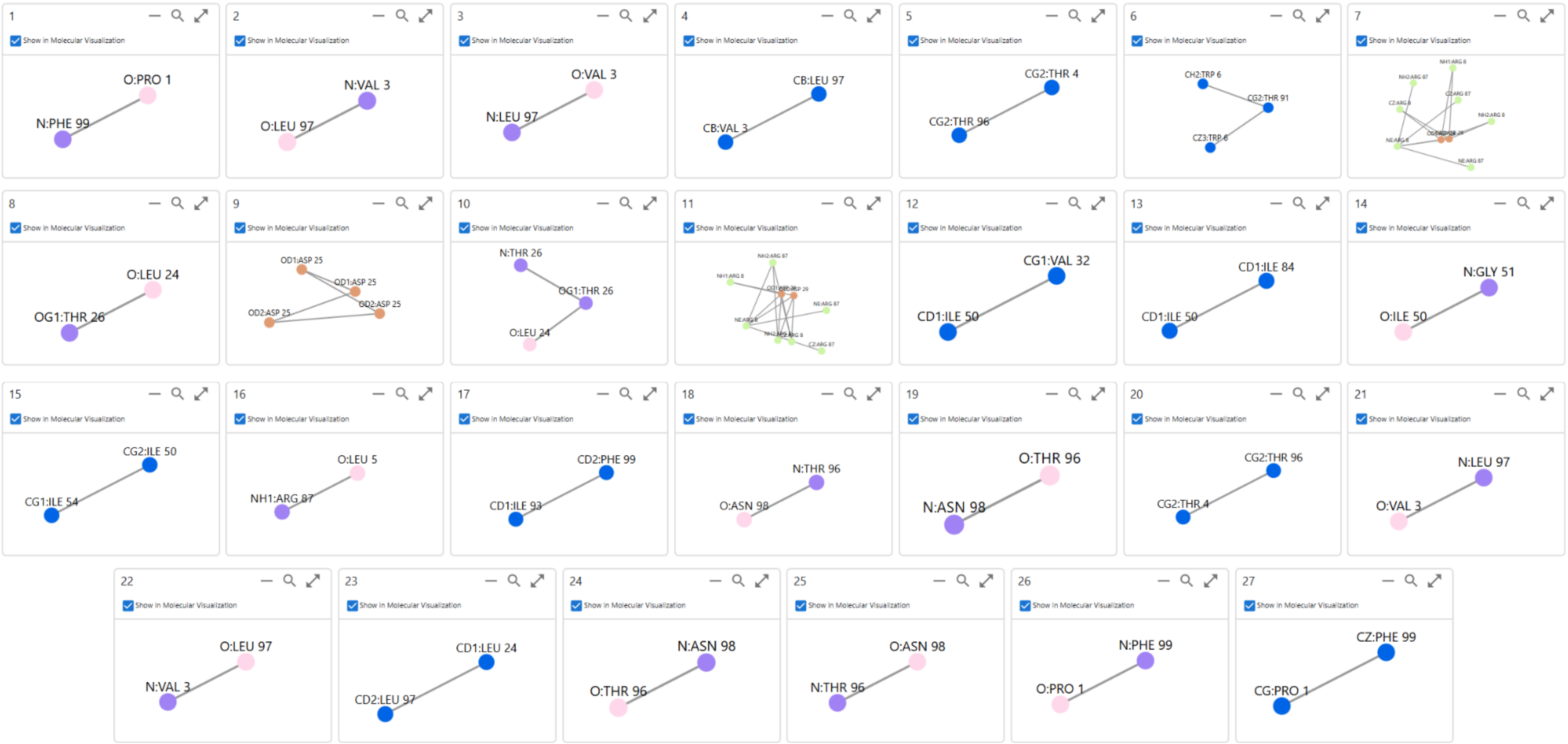


Set of 27 interaction graphs produced by PRIorI, each representing a specific pairwise interaction detected between residues of the HIV-1 protease and the bound inhibitor in the 1HVR structure. Each panel highlights the interacting atoms, interaction geometry, and spatial orientation relevant to the ligand–protein interaction network.

**Table 1 - Interaction profile between residues of the HIV-1 protease–ligand complex.**

| Index | Graph Number | Interaction Type | Atom 1 | Atom 2 | Distance |
|---|---|---|---|---|---|
| 1 | 1 | Hydrogen Bond | PRO-O-1-A (Acceptor) | PHE-N-99-B (Donor) | 2.89 Å |
| 2 | 2 | Hydrogen Bond | VAL-N-3-A (Donor) | LEU-O-97-B (Acceptor) | 2.64 Å |
| 3 | 3 | Hydrogen Bond | VAL-O-3-A (Acceptor) | LEU-N-97-B (Donor) | 2.79 Å |
| 4 | 4 | Hydrophobic | LEU-CB-97-B (Hydrophobic) | VAL-CB-3-A (Hydrophobic) | 3.60 Å |
| 5 | 5 | Hydrophobic | THR-CG2-4-A (Hydrophobic) | THR-CG2-96-B (Hydrophobic) | 3.58 Å |
| 6 | 6 | Hydrophobic | THR-CG2-91-B (Hydrophobic) | TRP-CZ3-6-A (Hydrophobic) | 3.52 Å |
| 7 | 6 | Hydrophobic | THR-CG2-91-B (Hydrophobic) | TRP-CH2-6-A (Hydrophobic) | 3.48 Å |
| 8 | 7 | Salt Bridge | ARG-NH1-8-A (Positive) | ASP-OD2-29-B (Negative) | 5.36 Å |
| 9 | 7 | Salt Bridge | ARG-NH1-8-A (Positive) | ASP-OD1-29-B (Negative) | 5.21 Å |
| 10 | 7 | Repulsive | ARG-NE-8-A (Positive) | ARG-NH2-87-B (Positive) | 4.82 Å |
| 11 | 7 | Salt Bridge | ARG-NE-8-A (Positive) | ASP-OD2-29-B (Negative) | 3.74 Å |
| 12 | 7 | Repulsive | ARG-NE-8-A (Positive) | ARG-NE-87-B (Positive) | 5.67 Å |
| 13 | 7 | Repulsive | ARG-NE-8-A (Positive) | ARG-CZ-87-B (Positive) | 5.65 Å |
| 14 | 7 | Salt Bridge | ARG-NE-8-A (Positive) | ASP-OD1-29-B (Negative) | 3.61 Å |
| 15 | 7 | Salt Bridge | ASP-OD2-29-B | ARG-CZ-8-A | 4.04 Å |

| | | | | | |
|---|---|---|---|---|---|
| | | | (Negative) | (Positive) | |
| 16 | 7 | Salt Bridge | ASP-OD2-29-B (Negative) | ARG-NH2-8-A (Positive) | 3.42 Å |
| 17 | 7 | Salt Bridge | ARG-CZ-8-A (Positive) | ASP-OD1-29-B (Negative) | 3.98 Å |
| 18 | 7 | Salt Bridge | ASP-OD1-29-B (Negative) | ARG-NH2-8-A (Positive) | 3.86 Å |
| 19 | 8 | Hydrogen Bond | LEU-O-24-A (Acceptor) | THR-OG1-26-B (Donor) | 2.83 Å |
| 20 | 9 | Repulsive | ASP-OD1-25-A (Negative) | ASP-OD2-25-B (Negative) | 4.51 Å |
| 21 | 9 | Repulsive | ASP-OD1-25-A (Negative) | ASP-OD1-25-B (Negative) | 2.99 Å |
| 22 | 9 | Repulsive | ASP-OD2-25-B (Negative) | ASP-OD2-25-A (Negative) | 5.59 Å |
| 23 | 9 | Repulsive | ASP-OD1-25-B (Negative) | ASP-OD2-25-A (Negative) | 4.77 Å |
| 24 | 10 | Hydrogen Bond | THR-OG1-26-A (Donor/Acceptor) | LEU-O-24-B (Acceptor) | 2.70 Å |
| 25 | 10 | Hydrogen Bond | THR-OG1-26-A (Donor/Acceptor) | THR-N-26-B (Donor) | 2.92 Å |
| 26 | 11 | Salt Bridge | ASP-OD2-29-A (Negative/Acceptor) | ARG-NE-8-B (Positive) | 4.08 Å |
| 27 | 11 | Salt Bridge | ASP-OD2-29-A (Negative/Acceptor) | ARG-CZ-8-B (Positive) | 3.69 Å |
| 28 | 11 | Salt Bridge | ASP-OD2-29-A (Negative/Acceptor) | ARG-NH1-8-B (Positive) | 4.55 Å |
| 29 | 11 | Salt Bridge | ARG-NE-8-B (Positive) | ASP-OD1-29-A (Negative) | 3.45 Å |
| 30 | 11 | Repulsive | ARG-NE-8-B | ARG-NE-87-A | 5.90 Å |

| | | | | | |
|---|---|---|---|---|---|
| | | | (Positive) | (Positive) | |
| 31 | 11 | Repulsive | ARG-NE-8-B (Positive) | ARG-CZ-87-A (Positive) | 5.74 Å |
| 32 | 11 | Repulsive | ARG-NE-8-B (Positive) | ARG-NH2-87-A (Positive) | 4.87 Å |
| 33 | 11 | Salt Bridge | ASP-OD1-29-A (Negative) | ARG-CZ-8-B (Positive) | 3.25 Å |
| 34 | 11 | Salt Bridge | ASP-OD1-29-A (Negative) | ARG-NH1-8-B (Positive) | 3.91 Å |
| 35 | 11 | Salt Bridge | ASP-OD1-29-A (Negative) | ARG-NH2-8-B (Positive/Donor) | 3.89 Å |
| 36 | 11 | Repulsive | ARG-CZ-8-B (Positive) | ARG-NH2-87-A (Positive) | 5.46 Å |
| 37 | 11 | Repulsive | ARG-NH2-87-A (Positive) | ARG-NH2-8-B (Positive/Donor) | 5.89 Å |
| 38 | 12 | Hydrophobic | VAL-CG1-32-A (Hydrophobic) | ILE-CD1-50-B (Hydrophobic) | 3.70 Å |
| 39 | 12 | Hydrophobic | ILE-CD1-84-B (Hydrophobic) | ILE-CD1-50-A (Hydrophobic) | 3.62 Å |
| 40 | 14 | Hydrogen Bond | GLY-N-51-A (Donor) | ILE-O-50-B (Acceptor) | 2.92 Å |
| 41 | 15 | Hydrophobic | ILE-CG2-50-B (Hydrophobic) | ILE-CG1-54-A (Hydrophobic) | 3.79 Å |
| 42 | 16 | Hydrogen Bond | LEU-O-5-B (Acceptor) | ARG-NH1-87-A (Donor) | 2.91 Å |
| 43 | 17 | Hydrophobic | PHE-CD2-99-B (Hydrophobic) | ILE-CD1-93-A (Hydrophobic) | 3.62 Å |
| 44 | 18 | Hydrogen Bond | THR-N-96-A (Donor) | ASN-O-98-B (Acceptor) | 2.79 Å |
| 45 | 19 | Hydrogen Bond | THR-O-96-A (Acceptor) | ASN-N-98-B (Donor) | 2.87 Å |
| 46 | 20 | Hydrophobic | THR-CG2-96-A (Hydrophobic) | THR-CG2-4-B (Hydrophobic) | 3.65 Å |

| | | | | | |
|---|---|---|---|---|---|
| 47 | 21 | Hydrogen Bond | LEU-N-97-A (Donor) | VAL-O-3-B (Acceptor) | 2.89 Å |
| 48 | 22 | Hydrogen Bond | LEU-O-97-A (Acceptor) | VAL-N-3-B (Donor) | 2.65 Å |
| 49 | 23 | Hydrophobic | LEU-CD1-24-B (Hydrophobic) | LEU-CD2-97-A (Hydrophobic) | 3.79 Å |
| 50 | 24 | Hydrogen Bond | ASN-N-98-A (Donor) | THR-O-96-B (Acceptor) | 2.86 Å |
| 51 | 25 | Hydrogen Bond | ASN-O-98-A (Acceptor) | THR-N-96-B (Donor) | 2.71 Å |
| 52 | 26 | Hydrogen Bond | PHE-N-99-A (Donor) | PRO-O-1-B (Acceptor) | 2.89 Å |
| 53 | 27 | Hydrophobic | PHE-CZ-99-A (Hydrophobic) | PRO-CG-1-B (Hydrophobic) | 3.56 Å |

Overall, the results obtained through PRIorI confirmed the presence of a network of non-covalent interactions between the inhibitor and the functional residues of the HIV-1 protease.
